# Comparative genomic analysis revealed specific mutation pattern between human coronavirus SARS-CoV-2 and Bat-SARSr-CoV RaTG13

**DOI:** 10.1101/2020.02.27.969006

**Authors:** Longxian Lv, Gaolei Li, Jinhui Chen, Xinle Liang, Yudong Li

## Abstract

The novel coronavirus SARS-CoV-2 (2019-nCoV) is a member of the family coronaviridae and contains a single-stranded RNA genome with positive-polarity. To reveal the evolution mechanism of SARS-CoV-2 genome, we performed comprehensive genomic analysis with newly sequenced SARS-CoV-2 strains and 20 closely related coronavirus strains. Among 98 nucleotide mutations at 93 sites of the genome among different SARS-CoV-2 strains, 58 of them caused amino acid change, indicating a result of neutral evolution. However, the ratio of nucleotide substitutions to amino acid substitutions of spike gene (9.07) between SARS-CoV-2 WIV04 and Bat-SARSr-CoV RaTG13 was extensively higher than those from comparisons between other coronaviruses (range 1.29 - 4.81). The elevated synonymous mutations between SARS-CoV-2 and RaTG13, suggesting they underwent stronger purifying selection. Moreover, their nucleotide substitutions are enriched with T:C transition, which is consistent with the mutation signature caused by deactivity of RNA 3’-to-5’ exoribonuclease (ExoN). The codon usage was similar between SARS-CoV-2 and other strains in beta-coronavirus lineage B, suggesting it had small impact on the mutation pattern. In comparison of SARS-CoV-2 WIV04 with Bat-SARSr-CoV RaTG13, the ratios of non-synonymous to synonymous substitution rates (dN/dS) was the lowest among all performed comparisons, reconfirming the evolution of SARS-CoV-2 under stringent selective pressure. Moreover, some sites of spike protein might be subjected to positive selection. Therefore, our results will help understanding the evolutionary mechanisms contribute to viral pathogenicity and its adaptation with hosts.

## Introduction

SARS-CoV-2, also known as 2019-nCoV, is a novel coronavirus (CoVs) isolated from patients with pneumonia in China 2019. SARS-CoV-2 has a similar incubation period (median, 3.0 days) and a relatively lower fatality rate than SARS-CoV or MERS-CoV (1), but it is estimated that the reproductive number of SARS-CoV-2 is higher than that of SARS-CoV (2). What’s more, some laboratory-confirmed symptomatic cases are absent of apparent cough, fever or radiologic manifestations, making it difficult to timely and accurately find out all infected patients (3, 4). As of February 28, 2020, patients infected by SARS-CoV-2 have been diagnosed in more than 30 countries, and more than 78,000 confirmed cases and 2,700 deaths associate with SARS-CoV-2 infection are reported in China alone.

The genetic information of a virus is essential for not only its classification and traceability, but also its pathogenicity. At the whole genome level, the sequence identify of SARS-CoV-2 was 50% to MERS-CoV, 79% to SARS-CoV, 88% to two bat-derived SARS-like coronaviruses, Bat-SL-CoVZC45 and Bat-SL-CoVZXC21 (collected in 2018 in Zhoushan, China), and 96% to Bat-SARSr-CoV RaTG13 (collected in 2013 in Yunnan, China) (5). Each genome of all SARS-CoV-2 strains now submitted online contains nearly 29,900 nucleotides (nt), which are predicted with at least 14 open reading frames (ORFs) (5′ to 3′), such as *ORF1ab* (P, 21,291 nt), *spike* (*S*, 3,822 nt), *ORF3a* (8,28 nt), *envelope* (*E,* 228 nt), *membrane* (*M,* 669nt), *ORF8* (366 nt), and *nucleocapsid* (*N,*1,260 nt) (6). Among them, the spike gene encoded a glycoprotein that is crucial for determining host tropism and transmission capacity, and highly divergent when compared with that of bat-SARSr-CoV RaTG13 (93.1% nucleotide identity) (6, 7).

Generally, the rates of nucleotide substitution of RNA viruses are faster than their hosts, and this rapid evolution is mainly shaped by natural selection (mostly purifying selection) (8). Gene mutations such as nucleotide substitutions, deletions and insertions have been frequently reported when comparing SARS-CoV-2 with other viruses (5–7). In this work, we investigated the mutation pattern of SARS-CoV-2 by comprehensive comparative genomic analysis of the nonsynonymous/synonymous substitution, relative synonymous codon usage (RSCU) and selective pressure in order to explore their potential in evolution and function.

## Materials and Methods

### Sequence data

The SARS-CoV-2 reference genome Wuhan-Hu-1 (NC_045512) and WIV04 (MN996528) was downloaded from the GenBank database. Other newly sequenced 2019-nCoV genomes were downloaded from the Global Initiative on Sharing Avian Influenza Data (GISAID) database (https://www.gisaid.org/). Twenty-one closely related coronavirus complete genome sequences and their coding sequences (CDS) were downloaded from GenBank database (Supplementary Table S1).

### Phylogenetic analysis

Genome sequences were aligned using MUSCLE v3.8.31 (9), followed by manual adjustment using BioEdit v7.2.5. Phylogenetic analyses of complete genome were performed using maximum-likelihood method and general time-reversible model of nucleotide substitution with gamma-distributed rates among sites (GTR+G) in RAxML v8.1.21 (10). Support for the inferred relationships was evaluated by a bootstrap analysis with 1000 replicates and trees were rooted with the alpha-coronavirus lineage as an outgroup.

The coding sequences were translated and aligned using the MEGA-X program (11), and then codon-based sequence alignment was used for further analysis. Phylogenetic analyses of coding sequences were performed using MEGA-X software. The changes of amino acids or nucleotides for each CDS sequences were analyzed using in-house Perl script.

### Estimation of synonymous and non-synonymous substitution rates

The number of synonymous substitutions per synonymous site (dS), and the number of non-synonymous substitutions per non-synonymous site (dN), for each coding region were calculated using the Nei-Gojobori method (Jukes-Cantor) in PAML package (12).

The adaptive evolution server (http://www.datamonkey.org/) was used to identify eventual sites of positive selection. For this purpose, the following test has been used: mixed-effects model of evolution (MEME), which allows the distribution of dN/dS (ω) to vary from site to site and from branch to branch at a site (13). This test allowed us to infer episodic and pervasive positive selection at individual sites.

### Synonymous Codon Usage Analysis

To investigate the potential relative synonymous codon usage (RSCU) bias of the spike protein from SARS-CoV-2 and its closely related coronaviruses, the coding sequence (CDS) of spike protein in these coronaviruses were calculated with CodonW 1.4.4 (http://codonw.sourceforge.net/). The RSCU of human genes was retrieved from the Codon Usage Database (http://www.kazusa.or.jp/codon/). The potential relationships among these sequences were calculated with a squared Euclidean distance 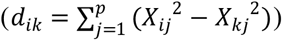. Besides RSCU, the effective number of codons (ENc) was used as a simple metrics to verify codon bias and to explore the source of virus.

### Statistical analysis

Statistical analyses were performed using the R statistical package (version 3.2.2). The Chi-squared test was used to compare any two data sets, and data were considered significantly different if the two-tailed p value was less than 0.05.

## Results and discussion

### The mutation pattern between SARS-CoV-2 and its closely related coronaviruses

The newly identified SARS-CoV-2 strain WIV04 genome sequence was closely related with Bat-SARSr-CoV RaTG13 and Bat-SL-CoVZC45, which was collected from host Rhinolophus affinis (5). Compared with RaTG13 genome, many nucleotide substitutions are observed, but there are only five small inserts and deletions (indels) mutations, and the largest insert segment in WIV04 genome was “CGGCGGGCACGT” sequence, which is located near the boundary of S1 and S2 regions of spike protein. Interestingly, only synonymous mutation are observed near this insertion sequence (Fig. 1, panel C). Compared with Bat-SL-CoVZC45 genome, this insert segment is detected as well, but non-synonymous mutations are also observed around it. Then, we further compared the proportion of synonymous mutations in spike gene between WIV04 and RaTG13 or Bat-SL-CoVZC45. The ratio of nucleotide substitutions (263 NT) to amino acid substitutions (29 AA) was 9.07 from WIV04 to RaTG13, which was significantly higher than the ratio (3.91, 864/221) from WIV04 to Bat-SL-CoVZC45 (p < 0.05) (Fig. 2). Moreover, we also observed that the synonymous mutations increased in whole genome level between WIV04 and RaTG13 than that between WIV04 and Bat-SL-CoVZC45.

**Figure 1.**
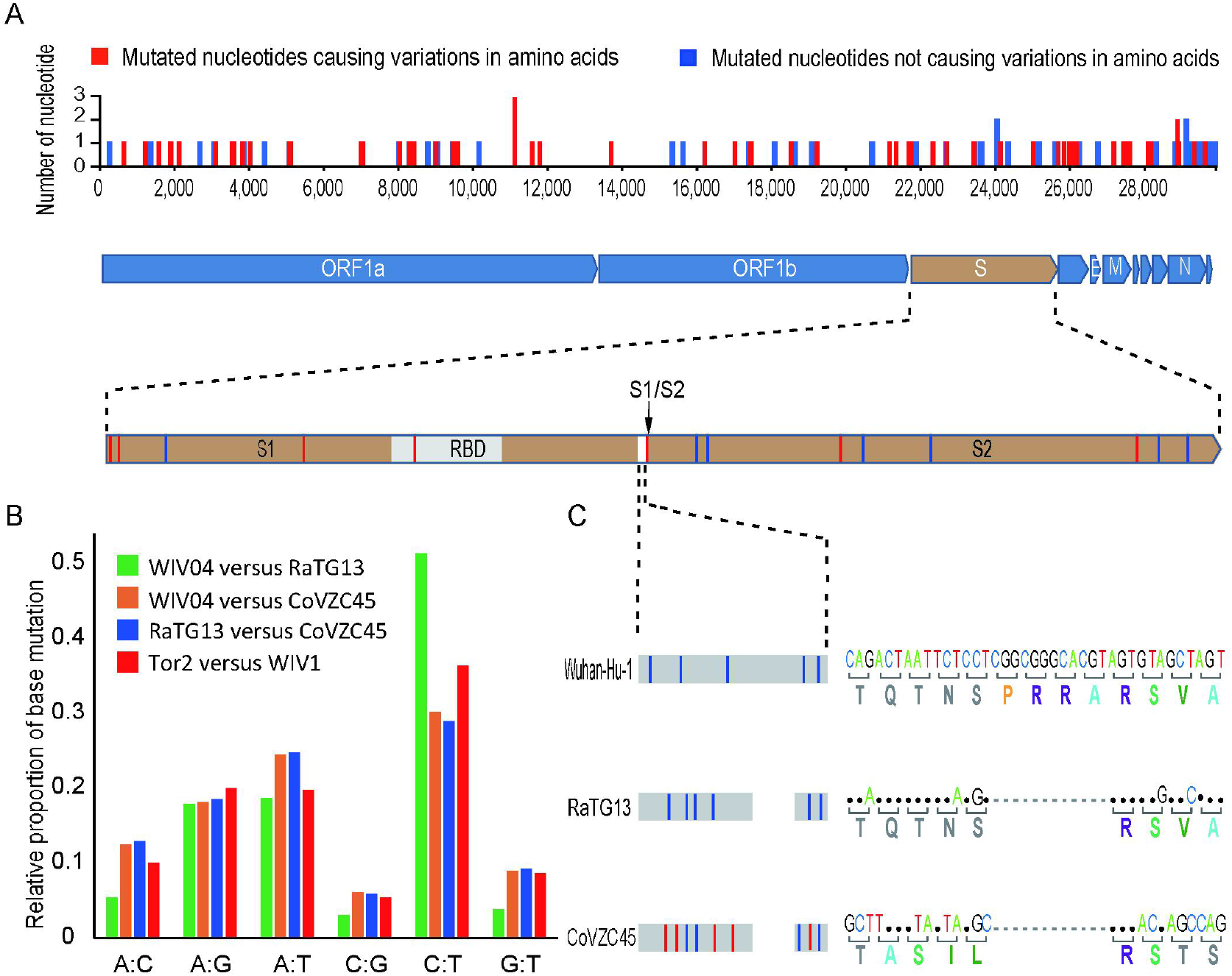
Mutation pattern in the coding region of SARS-CoV-2. (A) Point mutation distribution of 108 SARS-CoV-2 strains with genome available. The genetic diversity data of SARS-CoV-2 strains were collected from GISAID database (https://www.gisaid.org/epiflu-applications/next-sars-cov2-app/). The non-synonymous and synonymous mutations of SARS-CoV-2 viruses were highlighted with red or blue bar in the spike gene region. (B) Frequency of point mutations observed in spike gene. The frequency is the number of point mutations in each category (A:C, A:G, A:T, C:G, C:T, G:T) divided by total mutations. The “A:C” represents the nucloetide A was changed to C, or C changed to A. (C)The illustration of an insertion in spike sequence. The spike protein of SARS-CoV-2 was aligned against the most closely related SARS-like CoVs RaTG13 and Bat-SL-CoVCZ45.

**Figure 2.**
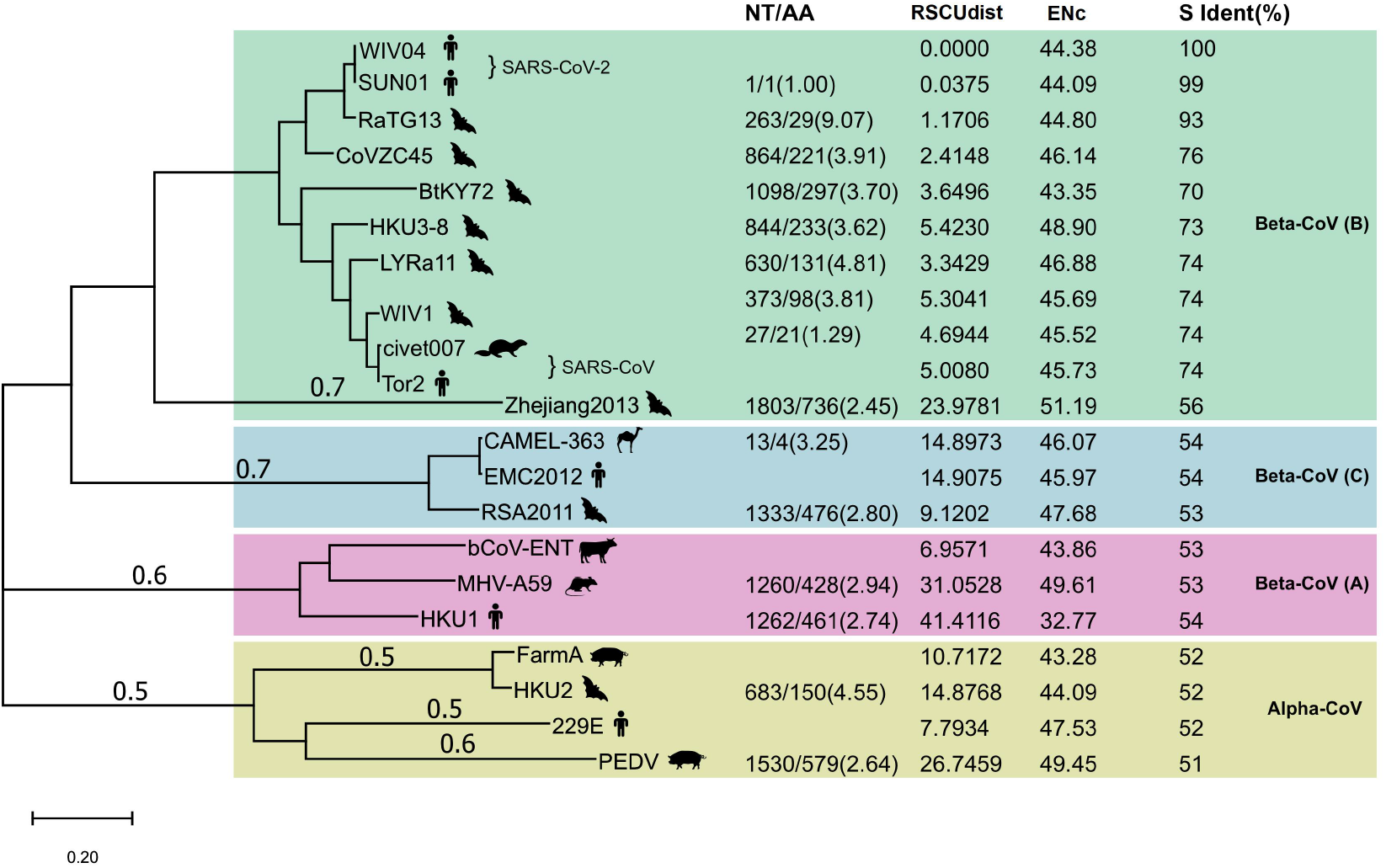
Maximum-likelihood phylogenetic tree of 21 coronavirus strains. The tree was built with full genome and rooted against alpha-CoV clade. The taxonomic name and NCBI accession number of each virus was list Supplemental table S1. NT: The number of nucleotide changes. AA: The number of amino acid changes. RSCUdist: the RSCU distance of spike protein; ENc: effective number of codons; S ident(%): the percent identity of S protein sequence between different coronaviruses. The NT/AA ratio between WIV04/RaTG13 was statistically significant higher (Chi-squared test, P<0.01) than that of other strain pairs.

Next, the detailed nucleotide changes of each comparison group were analyzed, and we found that the T-to-C (T:C) transition mutation was enriched in the nucleotide substitutions between WIV04 and RaTG13 (Fig. 1B). It’s reported that the CoVs lacking a 3’-to-5’ exoribonuclease (ExoN) accumulated 15 to 20 fold more A:G and U:C transition (14). What’s more, the RNA mutagen 5-fluorouracil (5-FU) treatment will also increase the U:C and A:G transitions. Consequently, this increased T:C mutation implies the ExoN of SARS-CoV-2 may be deactivated compared with that of RaTG13.

Furthermore, we checked whether the ratio of synonymous substitution to missense substitution is increased within SARS-CoV-2 strains. From December 2019 to February 2020, the genome sequences of 108 strains of SARS-CoV-2 virus have been submitted to GISAID database worldwide. Compared with the standard SARS-CoV-2 strain WIV04, total 98 point mutations were detected at 93 nucleotide sites of all SARS-CoV-2 strains with genome sequence available on Feb. 25 2020 (Fig. 1A). However, only 58 of these nucleotide mutations caused changes in amino acids. Among them, 15 nucleotide substitutions at 14 sites caused changes in 7 amino acids of the spike protein were observed. Consequently, the proportion of synonymous mutations (~40%) among all current reported SARS-CoV-2 strains is similar to that between WIV04 and Bat-SL-CoVZC45 (39.1%), but is dramatically less than that between WIV04 and RaTG13 (90.7%).

According to random drift hypothesis (15), these nucleotide mutations among different SARS-CoV-2 strains now available are still determined by neutral evolution. In short, there has no powerful factor to force SARS-CoV-2 to evolve in a certain direction by far. However, we should take strict precautions against the strong factors that may cause directional variation of SARS-CoV-2 both in natural environment and infection treatment.

### The mutation between SARS-CoV-2 and RaTG13 was unique across coronavirus species

To investigate whether the mutation pattern between SARS-CoV-2 and RaTG13 is unique across all coronavirus species, we further compared their alterations of nucleotides and amino acids in comparison to other representative coronaviruses. Phylogenetic analysis of SARS-CoV-2 and its 20 closely related coronaviruses formed four well supported clades (Fig. 2). The two SARS-CoV-2 strain WIV04 (from Wuhan) and SNU01 (from Korea patient) were clustered with SARS-CoV-related strains to form clade 1, belonging to beta-coronavirus lineage B. The MERS-CoV (EMC2012) from human, CAMEL-363 from camel and RSA2011 from bat formed clade 2, belonging to beta-coronavirus lineage C. The third beta-coronavirus lineage A was formed by the representive virus Bovine coronavirus (bCoV-ENT), Murine hepatitis virus (MHV-A59), and human coronavirus HKU1. The last clade was constituted of four representive alpha-coronaviruses. Notably, the sequence identities between closed related representative viruses, such as WIV04 and RaTG13, Tor2 and WIV1, SADS FarmA and HKU1 were nearly 95%.

Compared with Bat-SARSr-CoV RaTG13, there were 293 nucleotideswere altered while only 29 amino acids were altered in SARS-CoV-2; their ratio was as high as 9.07. However, this ratio detected in the comparisons of other human coronaviruses with their similar animal coronaviruses was less than 5.0. For example, the ratio of human SARS-CoV Tor2 to bat SARS-like CoVs LYRa11 was 4.81; that ratio of human MERS-CoV EMC2012 to bat SARS-like CoVs RSA2011 was 2.80; that ratio of human coronavirus 229E to bat coronavirus HKU2 was 4.55. These results indicate that the relative level of synonymous substitutions between human SARS-CoV-2 and its possible animal origin (RaTG13) is much higher than that between other human coronaviruses and their potential animal sources.

### Codon usage has small impact on the mutation pattern between SARS-CoV-2 and RaTG13

Different organisms, even different protein coding genes of the same species, have different frequency of codon usage (16). The RSCU bias will reveal the difference of the host source. We calculated the distance of RSCU between spike genes of 20 representative coronaviruses and SARS-COV-2. The codon usage difference (distance of RSCU) of between SARS-CoV-2 WIV04 and Bat-SARSr-CoV RaTG13 was 1.17, which was the lowest except for SNU01 (another strain of SARS-CoV-2), indicating that their codon preference was almost the same; the second lowest codon usage difference was 2.41 that was detected between SARS-CoV-2 WIV04 and Bat-SL-CoVZC45 (Fig. 2). The coronaviruses in same beta-CoV lineage B have a relatively close distance, except that of CoV Zhejiang2013. The codon usage difference of spike gene between human coronavirus HKU1 and bovine bCoV-ENT sequence have the largest difference, with a ratio of 41.41.

Codon usage bias in a gene can be effectively measured by determining the effective number of codons (ENc). Generally, the RNA viruses usually consist of high ENc values which help it in replication and host adaption with preferred codons. The lower ENc values represent high codon bias with low numbers of synonymous codons used for the amino acids, and a gene with strong codon usage bias may have an ENc value less than 35. The ENc values of WIV04 spike gene was 44.38 (Fig. 2), which is similar with those of RaTG13 and other bat coronavirus in Beta-CoVs lineage (B), indicating the high synonymous mutation was not determined by codon usage bias.

### The nucleotide substitutions between SARS-CoV-2 and RaTG13 was affected by stronger purifying selection

To infer whether the retention of synonymous mutations is supported or hindered by natural selection, we further studied non-synonymous substitution rate (dN) and synonymous substitution rate (dS) in spike gene (Table 1). Generally, positive (Darwinian) selection increases, but negative (purifying) selection decreases the ratios of non-synonymous to synonymous substitution rates (dN/dS). The results showed that both dN and dS of *S* gene of SARS-CoV-2 WIV04 versus Bat-SARr-CoV RaTG13 were the lowest among all typical coronaviruses, while those of SARS-CoV Tor2 versus bat SARS-like coronavirus WIV1 were the second lowest. When the ratio of dN to dS of spike gene is compared, all the tested dN/dS are less than 1, indicating that these non-synonymous mutations are harmful, and negative selection will reduce their retention speed. Among them, dN/dS of spike gene of SARS-CoV-2 WIV04 versus that of Bat-SARr-CoV RaTG13 was 0.04, which was the lowest among all comparisons, reconfirming that the rate of synonymous mutation was increased between WIV04 and RaTG13 strains. Moreover, dN/dS rate of polyprotein (ORF1ab) and nucleocapsid (N) genes were similar with that of spike gene (Table S2).

**Table 1.** Comparison of the evolutionary rate of spike protein and its RBD region between different coronavirus strains

| Coronavirus strains for |  | Spike |  |  | Receptor binding domain (RBD) |  |  |
| --- | --- | --- | --- | --- | --- | --- | --- |
| pair comparison |  | dN/dS | dN | dS | dN/dS | dN | dS |
| WIV04 | RaTG13* | 0.04 | 0.014 | 0.31 | 0.1165 | 0.064 | 0.5494 |
| WIV04 | CoVZC45 | 0.11 | 0.13 | 1.19 | 0.1067 | 0.2213 | 2.0751 |
| RaTG13 | CoVZC45 | 0.12 | 0.13 | 1.08 | 0.1067 | 0.2213 | 2.0751 |
| WIV04 | Tor2 | 0.11 | 0.17 | 1.50 | 0.095 | 0.028 | 0.2945 |
| Tor2 | WIV1* | 0.17 | 0.05 | 0.32 | 0.1592 | 0.1997 | 1.2545 |
| FarmA | HKU2* | 0.09 | 0.097 | 1.08 | 0.0835 | 0.1114 | 1.3342 |
| EMC2012 | RSA2011 | 0.21 | 0.29 | 1.38 | 0.4015 | 0.6464 | 1.61 |

\*The whole genome sequence identity between these paired strains was larger than 95%.

The spike (S) protein undergoes several drastic changes during virus infection. For instance, its large parts are cleaved during infection by cellular proteases and expose the receptors to activate viral attachment to the host (17). As the receptor binding domain (RBD) of spike protein was involved in interacting with human angiotensin-converting enzyme 2 (ACE2) protein, the RBD region was thought to be preferential targets of natural selection (18). Consistent with this hypothesis, our results showed that both dN and dS of RBD region are increased in comparison with whole spike gene region across all the virus pairs used in this study (Table 1). Notably, the dN/dS ratio of RBD region in SARS-CoV-2 WIV04 was dramatically increased in about 3 fold in comparison with full spike region. Consequently, these mutations may be subjected to Darwin’s choice and will become evidence of adaptive protein evolution.

To detect positive selection on the spike gene, the MEME analysis was performed. Significant (p < 0.05) pervasive episodic selection was detected in three sites (48th, 254th, 330th position using the reference sequence of WIV04). On the 254th position of the spike amino acid sequence, there is a histidine residue instead of a phenylalanine residue, whereas on the 330th amino acidic position in WIV04 sequence, there is a glutamine residue instead of an valine residue. The results described above strongly supporting the action of positive selection in spike gene during the recent evolution of SARS-CoV-2 and RaTG13. However, this result should be addressed with careful explain, because the RBD region of spike gene from SARS-CoV-2 was quit divergent with that of RaTG13 (Fig. S1), suggesting it might have originated from homologous recombination between RaTG13 and one yet-unknown coronavirus (19).

## Conclusions

Through comprehensive comparative analysis between SARS-CoV-2 and other coronaviruses, we found the synonymous mutations is dramatically elevated between SARS-CoV-2 and RaTG13 than that of other coronavirus strains, and the nucleotide mutations were enriched in T:C transition. The specific mutation pattern may caused by the loss function of RNA 3’-to-5’ exoribonuclease (ExoN). Moreover, as the SARS-CoV-2 was supposed to be originated from Bat-SARSr-CoV RaTG13, the increased synonymous substitution between SARS-CoV-2 and RaTG13 strain suggested the SARS-CoV-2 genome was under stronger negative (purifying) selection. We also detected some sites in spike protein was experiencing positive selection. These observations suggest that adaptive evolution might contribute to its host shifts. However, as the RNA mutagen (e.g. 5-FU) could induce the same mutation pattern, the mechanism of the mutation pattern observed between SARS-CoV-2 and RaTG13 should be further investigated in future work.

## Acknowledgments

We thank all the laboratories submitting the genome sequences of SARS-CoV-2 to GISAID or GenBank database for public research. Y. Li thank Prof. Wen-Hsiung Li to host him as visiting student to learn molecular evolution at his laboratory in Chicago University. This research was supported by the National Natural Science Foundation of China (31671836, 81570512), the National Key Research and Development Program of China (2018YFC2000500).

## Supplemental data

**Table S1.** The coronavirus genome sequences used in this study

| Strain name | Accession number | Host |
| --- | --- | --- |
| WIV04 | MN996528 | Human |
| SNU01 | MT039890 | Human |
| RaTG13 | MN996532 | Bat |
| CoVZC45 | MG772933 | Bat |
| Tor2 | AY274119 | Human |
| civet007 | AY572034 | Civet |
| WIV1 | KF367457 | Bat |
| LYRa11 | KF569996 | Bat |
| HKU3-8 | GQ153543 | Bat |
| BtKY72 | KY352407 | Bat |
| Zhejiang2013 | NC_025217 | Bat |
| EMC2012 | JX869059 | Human |
| CAMEL-363 | KJ713298 | Camel |
| RSA2011 | KC869678 | Bat |
| bCoV-ENT | NC_003045 | Bovine |
| MHV-A59 | NC_001846 | Mouse |
| HKU1 | NC_006577 | Bat |
| FarmA | MF094681 | Pig |
| HKU2 | NC_009988 | Bat |
| 229E | NC_002645 | Human |
| PEDV | NC_003436 | Pig |

**Table S2.** Comparison of the evolutionary rate of ORF1ab (P) and nucleocapsid (N) gene between different coronavirus strains

| Coronavirus strains for |  | P |  |  | N |  |  |
| --- | --- | --- | --- | --- | --- | --- | --- |
| pair comparison |  | dN/dS | dN | dS | dN/dS | dN | dS |
| WIV04 | RaTG13* | 0.0428 | 0.0052 | 0.1226 | 0.0453 | 0.0065 | 0.1433 |
| WIV04 | CoVZC45 | 0.1098 | 0.0354 | 0.3222 | 0.0494 | 0.0268 | 0.5422 |
| RaTG13 | CoVZC45 | 0.1109 | 0.0342 | 0.3085 | 0.0506 | 0.028 | 0.5544 |
| WIV04 | Tor2 | 0.0172 | 0.001 | 0.0607 | 0.0589 | 0.0077 | 0.1306 |
| Tor2 | WIV1* | 0.1458 | 0.0557 | 0.3819 | 0.0898 | 0.0968 | 1.078 |
| FarmA | HKU2* | 0.0929 | 0.025 | 0.2689 | 0.9776 | 0.3599 | 0.3682 |
| EMC2012 | RSA2011 | 0.1088 | 0.0542 | 0.4983 | 0.0785 | 0.0406 | 0.5167 |

\*The whole genome sequence identity between these paired strains was larger than 95%.

**Figure S1.**
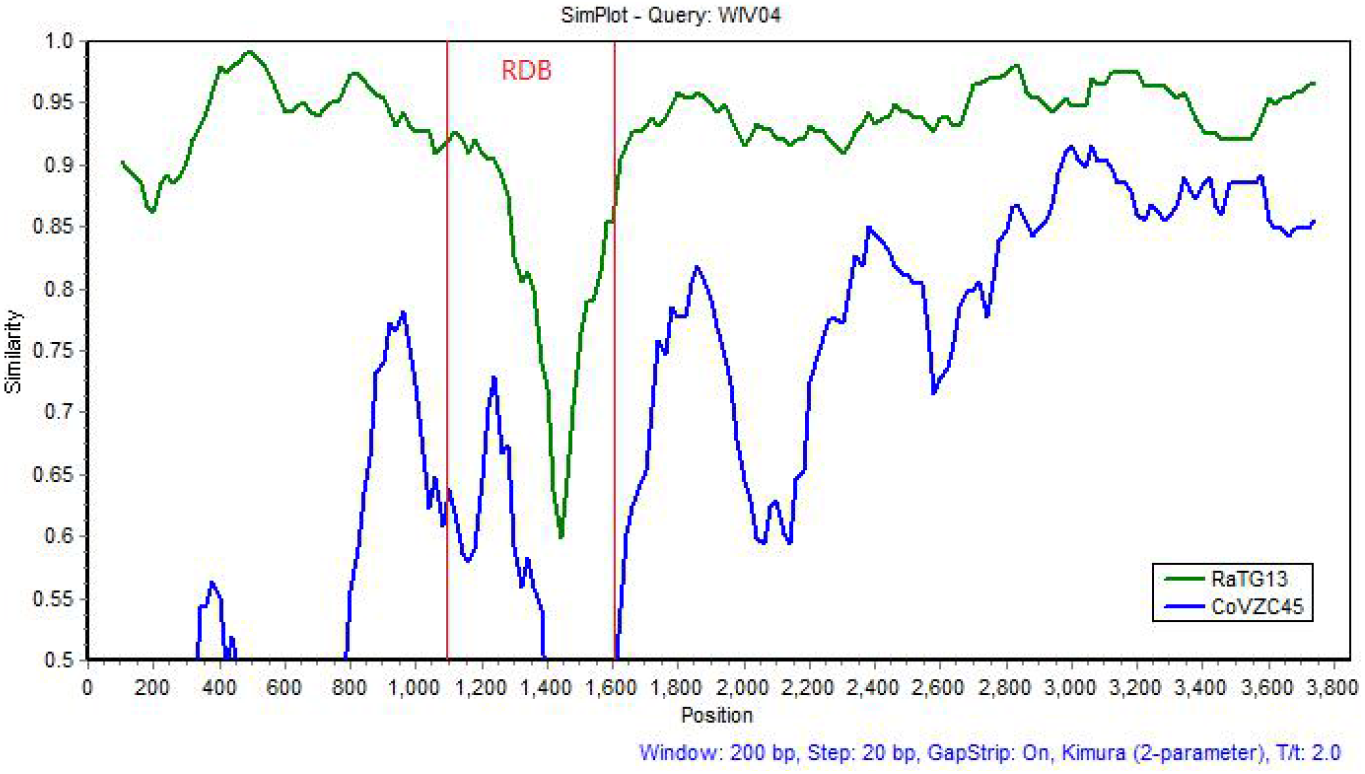
Sliding window analysis of changing patterns of spike gene sequence identity between SARS-CoV-2 (WIV04), RaTG13 and Bat-SL-CoVZC45. The position of RBD domain was shown in red lines.

